# The circadian clock regulates receptor-mediated immune responses to an herbivore-associated molecular pattern

**DOI:** 10.1101/2024.11.06.622352

**Authors:** Natalia Guayazán Palacios, Takato Imaizumi, Adam D. Steinbrenner

**Affiliations:** Department of Biology, University of Washington, Seattle, WA, USA

**Keywords:** HAMP, Inceptin, In11, immunity, legumes, time-of-day, circadian clock, circadian gating

## Abstract

Plants activate induced defenses through the recognition of molecular patterns. Like pathogen-associated molecular patterns (PAMPs), herbivore-associated molecular patterns (HAMPs) can be recognized by cell surface pattern recognition receptors leading to defensive transcriptional changes in host plants. Herbivore-induced defensive outputs are regulated by the circadian clock, but the underlying molecular mechanisms remain unknown. To investigate how the plant circadian clock regulates transcriptional reprogramming of a specific HAMP-induced pathway, we characterized the daytime and nighttime transcriptional response to caterpillar-derived In11 peptide, in the legume crop cowpea (*Vigna unguiculata*). Using diurnal and free-running conditions, we found that daytime In11 elicitation resulted in stronger late-induced gene expression than nighttime. Plants with a conditional arrhythmic phenotype in constant light (LL) conditions lost time-of-day dependent responses to In11 treatment, and this was associated with arrhythmic expression of circadian clock core transcription factor *Late Elongated Hypocotyl VuLHY1* and *VuLHY2*. Reporter assays with VuLHY homologs indicated that they interact with the promoter of daytime In11-induced *Kunitz Trypsin Inhibitor* (*VuKTI*) via a canonical and a polymorphic CCA1/LHY Binding Site (CBS), consistent with a mechanism of direct regulation by circadian clock transcription factors. This study improves our understanding of the time-dependent mechanisms that regulate herbivore-induced gene expression.

## Introduction

Plant survival is dictated by the plant’s ability to accurately perceive biotic threats and to activate effective defenses in a timely manner. Plants sense pests and pathogens through recognition of molecular patterns via cell-surface pattern recognition receptors (Ngou *et al*., 2022; Zhang *et al*., 2024). The specific interaction between a molecular pattern and a cognate receptor results in the activation of Pattern Triggered Immunity (PTI) to provide a first line of protection, a hallmark of which is the transcriptional reprogramming that accompanies metabolic and physiological changes associated with immunity (DeFalco & Zipfel, 2021). While gene expression changes and the mechanisms regulating them in response to Pathogen Associated Molecular Patterns (PAMPs) are well documented (Li *et al*., 2016; Bjornson *et al*., 2021), our understanding of these processes in response to Herbivore Associated Molecular Patterns (HAMPs) is nascent. Although many HAMPs have been identified (Snoeck *et al*., 2022b), detailed transcriptional responses to specific elicitor molecules have only been described for two HAMPs present in lepidopteran oral secretions: the fatty acid-amino acid conjugate (FAC) C18:3-Glu in *Nicotiana* species (Zhou *et al*., 2016), and the peptide Inceptin 11 (In11) active on cowpea, *Vigna unguiculata* (Steinbrenner *et al*., 2022). While FAC receptors are not yet fully elucidated (Poretsky *et al*., 2020), In11 is recognized only in select legume species due to a legume-specific Inceptin Receptor (INR), a leucine-rich repeat receptor in the Receptor Like Protein (RLP) family (Steinbrenner *et al*., 2020; Snoeck *et al*., 2022a). Since In11-INR is the only HAMP-receptor pair characterized in molecular detail it serves as a model for studying herbivore-specific immune pathways (Steinbrenner *et al*., 2022)

In11 elicitation results in a well-characterized set of defensive outputs driven by amplified and accelerated expression of wound-induced genes, as well as In11-specific gene expression (Steinbrenner *et al*., 2022). As a result of rapid transcriptional reprogramming in response to In11, both direct and indirect induced defenses are accumulated to increase resistance to herbivores (Schmelz *et al*., 2006). Induced defense responses include production of defense-related phytohormones, specialized anti-nutritive proteins and metabolites, and volatile-mediated attraction of beneficial insects (Erb & Reymond, 2019). HAMP-induced regulation of these responses is thought to be a mechanism to reduce costs by effectively allocating defenses to times and tissues when and where they are needed (Karban, 2011).

Like many other biological processes, plant immunity is regulated by the circadian clock (Lu *et al*., 2017). The core circadian clock components CIRCADIAN CLOCK ASSOCIATED 1 (CCA1), LATE ELONGATED HYPOCOTYL (LHY) and LUX ARRYTHMO (LUX) are Myb-like transcription factors that participate in the rhythmic accumulation of defensive hormones jasmonic (JA) and salicylic acid (SA) (Goodspeed *et al*., 2012), and resistance genes such as *RECOGNITION OF PERONOSPORA PARASITICA 4* (*RPP4*) (Wang *et al*., 2011) in anticipation to herbivore and pathogen attack, respectively. LUX is also involved in modulating PTI responses in a time-of-day-dependent manner through gated accumulation of ROS and expression of the bacterial marker gene *FLG22-INDUCED RECEPTOR-LIKE KINASE 1* (*FRK1*) in response to flagellin 22 (flg22), a bacterial PAMP, in the early morning (Korneli *et al*., 2014). Whether HAMP-induced transcriptional changes are also time-of-day dependent and if they are modulated by the plant circadian clock remains unknown.

Modulation of gene expression is directly regulated by clock transcription factors. For example, CCA1 and LHY bind the cis-regulatory elements CCA1 Binding Site (CBS) (Wang *et al*., 1997) and Evening Element (EE) (Harmer *et al*., 2000) to regulate target gene expression via repression and/or activation (Wang *et al*., 2011; Nagel *et al*., 2015). CBS and EE cis-elements have been found in the promoter of rhythmically expressed bacterial resistance genes (Wang *et al*., 2011), and the herbivore-induced *Ocimene Synthase* (*PlOS)* in lima bean (*Phaseolus lunatus*), a transcript rhythmically accumulated in response to herbivore feeding and regulated by light and JA (Arimura *et al*., 2008). However, studies of herbivory and HAMPs have not yet measured whether direct regulation by clock transcription factors extends to genome-wide changes in herbivore-induced gene expression.

Here we present a detailed exploration of the temporal induced response to HAMP In11 and provide a molecular link between the In11-induced immune responses and the plant circadian clock. We characterized the global early and late transcriptional changes induced by daytime and nighttime In11 treatment and identified a daytime-induced antiherbivore-related *Kunitz Trypsin Inhibitor* (*KTI*) gene with CBS elements in the promoter region. Using plants with a conditional arrhythmic phenotype under constant light we tested if the daytime *KTI* induction in response to In11 required a functioning clock and found that the misexpression of *VuLHY* homologs under constant light (LL) correlated with lack of repression of *KTI* during nighttime. Furthermore, we show that transient overexpression of cowpea LHY proteins in *Nicotiana benthamiana* modulates the activity of the *Vu*KTI promoter in a CBS-dependent manner. We propose that VuLHY transcriptionally gates In11-induced gene expression at night to ensure that specific defenses are most strongly produced in response to herbivorous threats in daytime.

## Materials and Methods

### Plant materials and growth conditions

Cowpea (*Vigna unguiculata*) accession IT97K-499-35 was used in all the experiments. For planting, seeds were surface sterilized with 70% ethanol for 2 minutes, followed by two washes with sterilized water. Seeds were sown on sunshine potting mix No.5 and placed in a growth chamber (Conviron PGW-40) at 26 C, 70% relative humidity (RH), 500 µmol/m^2^sec light intensity, and 12 h light/dark (LD) cycle for 14 days. Details specific to diurnal and LL experiments are provided in the following sections.

### In11 treatment under diurnal and constant light conditions

Inceptin 11 (In11) peptide (ICDINGVCVDA) was synthesized (Genscript Inc.) and dissolved in water. We lightly wounded the middle leaflet of the first fully extended trifoliate on 14-day-old cowpea plants using a new razor blade to remove the cuticle (1 cm^2^ per wound). We made four wounds, two on each side of the main vein of the adaxial side of the leaflet, and equally distributed 20 µL of either water or 1 µM In11 with a pipette tip.

For the diurnal experiment, we applied the In11 and water treatments 4 h after the lights came on (daytime, Zeitgeber time 4: ZT4) or 4 h after the lights went off (nighttime, Zeitgeber time 16: ZT16) in the growth chamber. We collected samples 1 h (ZT5 and ZT17) and 6 h (ZT10 and ZT22) after treatment, along with untreated controls. For the constant light (LL) experiment, we transferred LD-grown cowpea plants (see above) 10, 11, 12 or 13 days after germination to a separate growth chamber under LL. On day 14, we treated all plants with In11 or water at subjective daytime (time in LL: 4, 28, 52 and 76 h) or subjective nighttime (time in LL: 16, 40, 64 and 88 h). We then collected samples from independent plants 6 h after treatment (time in LL daytime 10, 34, 58, 82 h and nighttime 22, 46, 70, 94 h), along with untreated controls.

### RNA extraction, qRT-PCR and transcriptomics

In all experiments we collected samples as follows: two leaf discs were taken from the treated leaflet (one proximal and one distal) using a 0.6 cm^2^ leaf punch, placed in a 1.5 mL tube containing a metal bead, frozen in liquid nitrogen, and stored at -80 C. Prior to RNA extraction, the samples were ground using a mixer mill (Retsch MM400).

For qPCR, we extracted total RNA using the Trizol (Invitrogen) method. We performed quality control of the RNA by NanoDrop1000 and gel electrophoresis and 1 µg of RNA was used to synthesize cDNA using the SuperScript IV RT Kit (Thermo). We used the Power SYBR™ Green PCR Master Mix for amplification and quantification in a CFX Connect Real-Time System (Bio-Rad). We calculated relative gene expression by using the 2^-ΔΔCt^ method and *Ubiquitin (UBQ)* (*Vigun07g244400*) as an expression control (Table S1).

For RNAseq, we extracted total RNA using the NucleoSpin Plant RNA kit (Macherey-Nagel Inc.) and performed quality control as explained above. We further treated the RNA using the TURBO DNA-free kit (Invitrogen) as DNA was still present in the samples. The extracted RNA was used to generate paired-end Illumina 2×150 bp strand-specific libraries with polyA selection that were sequenced in a HiSeq2500 (Azenta). For gene expression analyses, we mapped the reads to the cowpea genome (Liang *et al*., 2024a) *Vigna unguiculata* v1.2 available in Phytozome13 (Goodstein *et al*., 2012) and used the –quantMode in STAR to quantify them (Dobin *et al*., 2013), and then performed differential gene expression analyses using DESeq2 (Love *et al*., 2014) implemented in R.

### Motif search

We searched known CCA1 and LHY binding sites in the promoters from all genes in the cowpea genome (*Vigna unguiculata* v1.2). We retrieved the 1.5 kb region upstream of the start codon from all genes using a custom Python script and the annotation file, and then used the Find Individual Motif Occurrences (FIMO) (Grant *et al*., 2011) online tool to find the 8-mer “AA**MW**ATCT”, where M was Adenine (A)/Cytosine (C) and W was Adenine (A)/Thymine (T). We selected this motif because it represented all possible CCA1 binding sites (CBS, CBS-A: AA**AA**ATCT and CBS-B: AA**CA**ATCT) and evening element (EE, AA**AT**ATCT) sequences (Table S2).

### Phylogenetic analysis

We used Arabidopsis (*Arabidopsis thaliana*) CCA1 (AT2G46830.1) and LHY (AT1G01060.1) as queries to retrieve sequences from cowpea, common bean (*Phaseolus vulgaris*), lima bean (*Phaseolus lunatus*), soybean (*Glycine max*), and Medicago (*Medicago truncatula*) genomes available in Phytozome13 using tblastn. We retained the top 30 similar sequences and aligned them using MAFFT v7.48 (Katoh *et al*., 2002) with default parameters. We constructed a phylogenetic tree using RAxML v8 (Stamatakis, 2014), used FigTree (http://tree.bio.ed.ac.uk/software/figtree/) to root and visualize the tree, and manipulated the image in Adobe Illustrator 2024.

### Molecular Cloning of *VuLHY* homologs and *pKTI* promoter

For plant protein expression, the full length coding sequences of cowpea *VuLHY1* (*Vigun10g1533300*) and *VuLHY2* (*Vigun09g004100*) were amplified from a 5’ RACE cDNA library using the Q5 High-Fidelity DNA polymerase (NEB) and specific primers (Table S1). The PCR products were cloned into pENTR D-TOPO (Thermo Fisher) and recombined into pB7WG2 for plant expression using the Gateway LR Clonase II (Invitrogen) (Karimi *et al*., 2002).

For the luciferase reporter assays, the promoter sequence of Kunitz Trypsin Inhibitor (*pKTI*) *Vigun05g143300* (region from the start codon up to 1 kb upstream) was amplified from genomic DNA using DreamTaq DNA polymerase (Thermo Fisher) and specific primers (Table S1) designed against the *V.unguiculata* v1.2 genome. The resulting fragment was cloned into pENTR 5’-TOPO (Thermo Fisher). Mutant versions of the promoter were generated from the wild type (WT) clone using the Q5 Site-Directed Mutagenesis Kit (NEB) and mutagenic primers (Table S1). WT and mutant promoters were re-amplified from the pENTR 5’-TOPO clones using primers with added BpiI restriction enzyme recognition sites and overhangs compatible with Mo-Clo and cloned into a level 0 cloning vector (pICH41295, Addgene plasmid # 47997). The reporter construct was assembled into a customized pGreenII (Hellens *et al*., 2000) with BsaI insertion site by combining appropriate ratios of the promoter, luciferase CDS (pICSL80001, Addgene plasmid # 50326) and *ocs* terminator (pICH41432, Addgene plasmid # 50343) following the recommended XL ligation protocol (Weber *et al*., 2011; Engler *et al*., 2014)

### Transient luciferase assay in *Nicotiana benthamiana*

To test the *in-planta* interaction between *pKTI* and the VuLHY proteins, we individually transformed pGII reporter and pB7WG2 effector constructs into Agrobacterium (*Agrobacterium tumefaciens* GV3101). Cultures were resuspended in infiltration media (10 mM MES pH 5.6, 150 μM Acetosyringone, 10mM MgCl_2_) and incubated for 3 h. For co-infiltrations, we prepared the appropriate combinations of the reporter and effector. To account for transformation efficiency and to enhance protein expression, we included *35S:Renilla* (final OD_600_=0.1) and the tomato stunt bushy virus silencing-suppressor p19 (final OD_600_=0.1) plasmid in all our assays. The youngest fully expanded leaf on a 6-week-old *Nicotiana benthamiana* plant was infiltrated with the mixture using a needleless syringe at Zeitgeber time 6 (ZT6). All plants were entrained to 12 h light/12 h dark cycles and, after 74 h of incubation at ZT8, we collected leaf punches and immediately froze them in liquid nitrogen. We prepared and analyzed the samples with the Dual-Luciferase Assay System (Promega) according to manufacturer instructions. We measured the activities of firefly (LUC) and Renilla (REN) luciferases using a multi-mode plate reader (Tecan Spark) and calculated the LUC/REN ratio for each reporter - effector combination.

### Statistical analysis

All statistical analyses were conducted using R version 4.3.2, and the significance level was set to =0.05. Extreme outliers were identified using the *identify_outliers* function from the rstatix (https://CRAN.R-project.org/package=rstatix) package and removed from the qPCR data. A normal distribution of the residuals was confirmed for all data using the *Shapiro.test* function, and transformations were applied when appropriate to fulfill the assumption of normality for ANOVA and t-test. ANOVA and Tukey post hoc test were used to identify significantly different means of gene expression across treatments, and a two-sided t-test was used to determine significantly different means of promoter activity in the presence of a VuLHY homolog vs an empty vector (EV).

## Results

### In11-induced transcriptional responses are dependent on the time-of-the-day

To examine the contribution of time-of-day on the transcriptional response to a specific HAMP (In11) via a known receptor (INR), we treated cowpea plants by scratch-wounding and adding water (w + H_2_O) or In11 (w + In11) at different times of the day (Fig. **1a**). We identified differentially expressed genes (DEGs) by comparing transcriptomes of w + H_2_O vs undamaged (i.e. the effect of wounding), and w + In11 vs. w + H_2_O treatment (i.e. the additional effect of In11-induced responses) at the corresponding time of the day (|log_2_ fold change (FC)| ≥1 and P_adj_ <0.05) (Table S3). A Principal component (PC) analysis across all samples confirmed consistent biological replicates and the effect of the treatments. The largest changes are attributed to damage and time, with clear separation of the early 1 h wound responses, late 6 h wound responses, and undamaged plants. Within those groups, there are also differences in the response to In11 depending on the time-of-day at which the treatment was applied (Fig. **S1a**).

**Figure 1.**
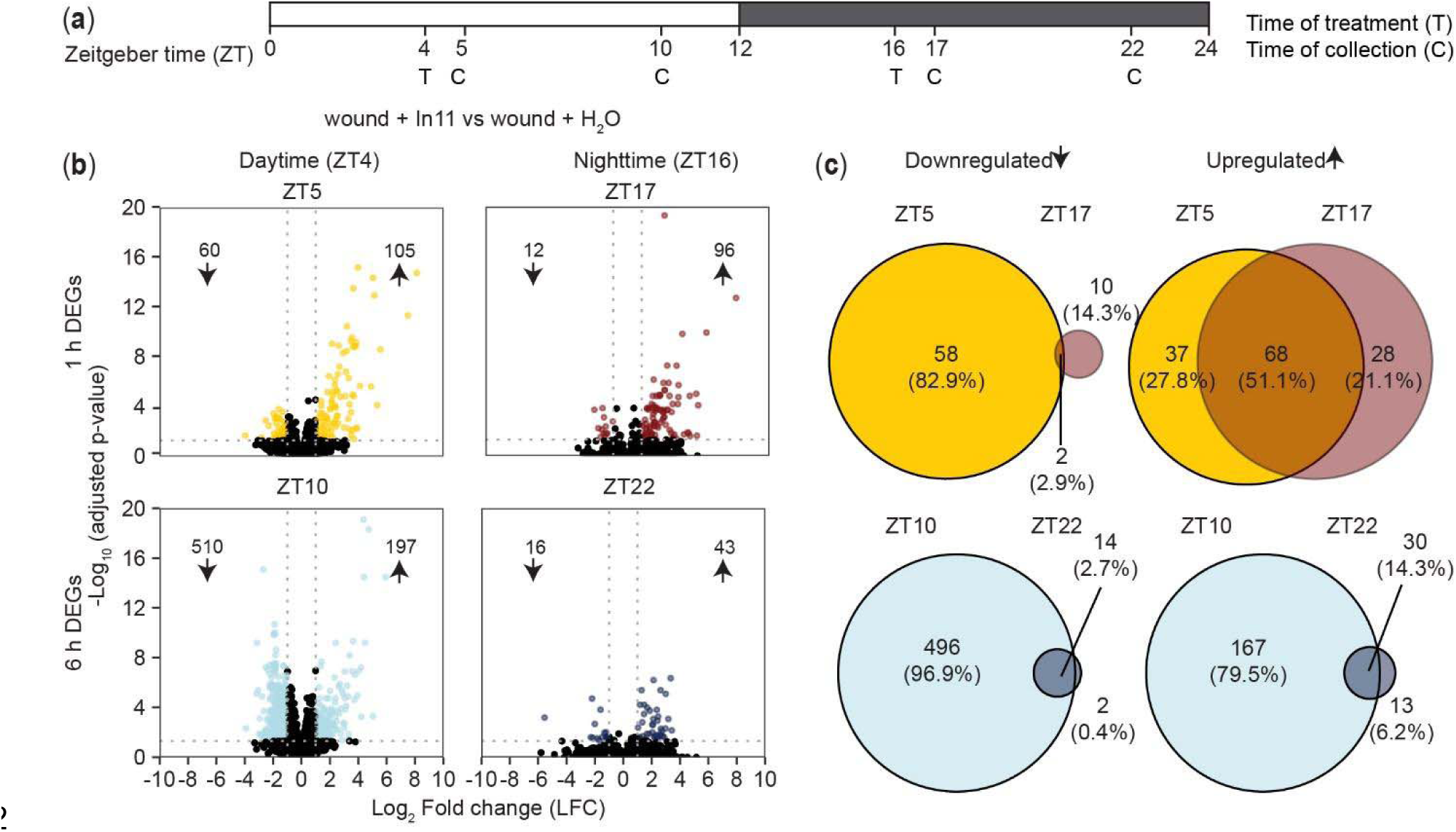
In11 induced responses are time-of-day dependent. (a) Experimental design for RNA-seq. 14-day old cowpea plants grown under diurnal conditions (light/dark, LD) were treated (T) with wound + H_2_O or wound + In11 at daytime (ZT4) or nighttime (ZT16), and samples were collected (C) 1h (ZT5 and ZT17) and 6h (ZT10 and ZT22) after treatment (n = 4 individual plants as biological replicates). (b) Volcano plots displaying the number of In11 down (↓) and upregulated (↑) genes (Log_2_ Fold Change|LFC| ≥ 1 relative to wound + H_2_O and p_adj_ < 0.05) 1h and 6h after treatment. (c) Venn diagram indicating the number of shared and unique differentially expressed genes (DEGs) 1h (ZT5 vs ZT17) and 6h (ZT10 vs ZT22) after daytime or nighttime treatment.

We found a time-of-day dependent response to In11 where daytime treatment resulted in a larger number of transcriptional changes than nighttime treatment (Fig. **1b,c**). While this was true for both 1 h and 6 h responses, time-of-day dependence of the In11 response was particularly striking 6 h after treatment as there were 707 DEGs at ZT10 (light blue, 510 down and 197 up) but only 59 at the corresponding nighttime timepoint ZT22 (dark blue, 16 down and 43 up). Furthermore, most DEGs were unique to ZT10 with only 44 shared with ZT22. A hierarchical clustering analysis of all In11 DEGs further supported the unique In11-induced transcriptional program at ZT10, and revealed that In11-induced nighttime responses were more similar to wounding alone because w+In11 ZT22 samples clustered most closely with day- and nighttime w+H_2_O plants, rather than with w+In11 at ZT10 (Fig. **S1b**). Interestingly, these patterns for In11-regulated genes did not hold for the broader set of 15,842 genes affected by wounding (Fig. **S2a**). In contrast to nearly complete time-of-day dependence of In11-induced downregulation at the 6 h timepoint, wound-induced downregulation of genes was intact at night (ZT22), and affected an even larger number of genes than in daytime (ZT10); nevertheless, nearly 50% of the up and downregulated genes were shared between daytime and nighttime (Fig. **S2b**). Together these results indicate that In11 modulates the wound response in a time-of-day dependent manner.

We compared the types of genes in the daytime and nighttime DEGs to identify shared and unique processes modulated by the recognition of In11 at different times of the day. Gene Ontology (GO) analysis revealed that daytime upregulated genes were significantly enriched for molecular functions related to antiherbivore defense such as lipid biosynthesis and metabolism, acyltransferase activity, protease binding and terpene synthase activity, while the downregulated DEGs were enriched for photosynthesis (Table S4).

### Direct and indirect antiherbivore defenses may be directly regulated by the circadian clock

Given the time-of-day dependent response to In11 we hypothesized that the circadian clock could be directly modulating gene expression; specifically, that cowpea homologs of the transcription factors CCA1 or LHY were directly repressing gene expression at nighttime via canonical cis elements CCA1 binding site CBS (Wang et al., 1997) and evening element (EE) (Harmer et al., 2000). To find evidence for direct CCA1/LHY regulation in specific promoter sequences, we calculated the LFC difference (LFC_diff_) between daytime and nighttime treatments for all In11-induced DEGs, calculated by comparing ZT5 vs ZT17 for 1 hr differences, and ZT10 vs ZT22 for 6 hr differences (Fig. 2a). We then annotated 21,768 total CBS and EE sequences in their promoters using FIMO (Fig. **S3a**). We focused on the 6 h comparison because of the strong effect of time-of-day, and found that 326 out of 722 unique DEGs (45.1%) across ZT10 and ZT22 had at least one of the cis elements (purple dots, Fig. 2a), and of those 104 had LFC_diff_ ≥1 (Table S5). This was a similar proportion of CBS and EE to the entire genome 14,861 out of 31,948 genes with at least one element (46.5%). We focused on candidate targets with defense-related functions for further analysis (Fig. **2a**), and found that genes with functions in indirect and direct defenses, such as terpene synthases (TPS) and Kunitz Trypsin Inhibitors (KTI), respectively, contain CBS and/or EE sites in promoters, suggesting they are a target of CCA1/LHY (Fig. **2b**). Given that the accumulation of certain induced indirect defenses in response to herbivory is known to be time-of-day dependent (Arimura *et al*., 2008), we selected *VuKTI* (*Vigun05g143300*), encoding a direct defense, as a marker gene. We confirmed by qPCR that *VuKTI* was significantly more induced 6 h after daytime application of In11 (ZT10), but not 6 h after nighttime treatment (ZT22) compared to wounding alone (Fig. **2c**).

**Figure 2.**
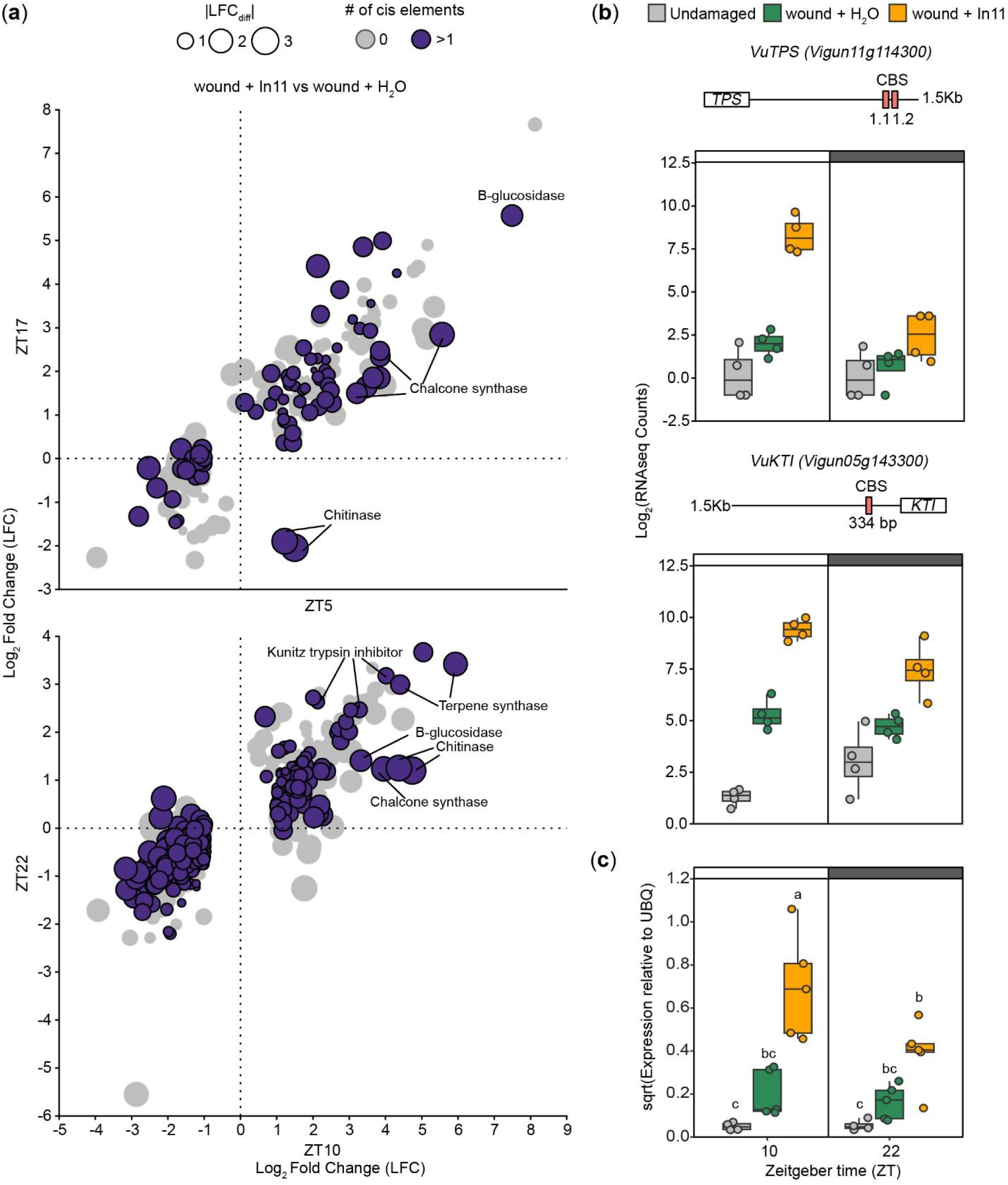
Circadian clock related cis elements CBS and EE are present in the promoters of time-of-day dependent In11-induced defense genes. (a) Scatter plot showing the Log_2_ Fold Change (LFC) value of In11 DEGs (wound + In11 vs wound + H_2_O) at 1h (ZT5 and ZT17) and 6h (ZT10 and ZT22) after treatment, and the absence/presence (gray/purple circles) of CBS or EE in their promoter (1.5 kb upstream start codon). The absolute value of the LFC difference (|LFC_diff_|) is represented by the size of the circles, and selected defense-related genes are indicated. (b-c) Promoter structure and expression pattern of a Terpene Synthase (*VuTPS*) and Kunitz Trypsin Inhibitor (*VuKTI*) 6h (ZT10 and ZT22) after daytime and nighttime treatment. according to (b) RNAseq data (b) and qPCR data (c) are shown. Different letters indicate significant differences determined by two-way ANOVA followed by Tukey’s Honest Significant Difference test (HSD) (n= 4-5 biological replicates, p-value < 0.05). Independent plants were sampled at each treatment - time combination.

### Expression of *VuLHY* homologs is disrupted by wounding and constant light in cowpea

To further investigate circadian clock modulation of In11 induced defenses, we first identified LHY in cowpea and profiled its expression pattern under various conditions. We identified two homologs *VuLHY1* (*Vigun10g153300*) and *VuLHY2* (*Vigun09g004100*) (Fig. **3a**) and determined that their transcripts have rhythmic expression that peaks at dawn (ZT4) in LD in undamaged samples (gray lines) according to our transcriptomics (Fig. **3b**) and independent qPCR data (Fig. **3c**); although *VuLHY1* expression was stronger than *VuLHY2*. Furthermore, both genes were significantly downregulated (Table S1) in response to wounding at nighttime ( ZT22), without further effect of In11. Together, these results suggest a reciprocal regulation of the circadian clock by wounding in cowpea.

**Figure 3.**
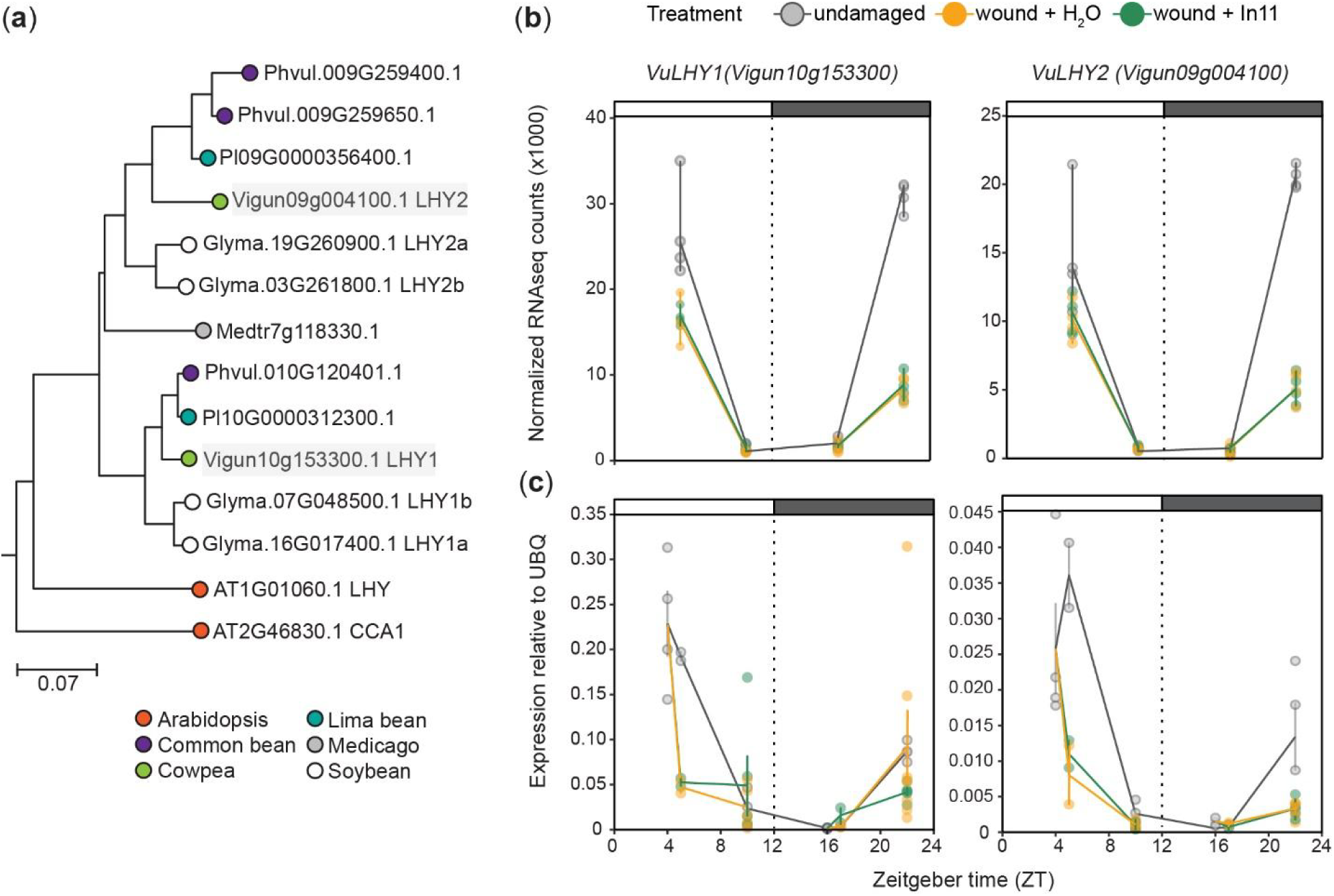
Cowpea Late Elongated Hypocotyl (*VuLHY*) homologs show typical cycling patterns and are downregulated by wounding. (a) Maximum likelihood phylogenetic tree showing 14 *LHY* homologs from five legume species and Arabidopsis. Cowpea homologs *VuLHY1* and *VuLHY2* are highlighted in gray boxes. The scale bar indicates branch length as the mean number substitutions per site. Diurnal expression pattern of *VuLHY*1 and *VuLHY*2 according to (b) RNAseq (n = 4) and (c) qPCR data (n = 3-4 biological replicates). Samples were collected at ZT4, ZT5, ZT16, ZT17 and ZT22 from undamaged plants (gray), and 1(ZT5, ZT17) and 6 h (ZT10, ZT22) after daytime (ZT4) or nighttime (ZT16) wound + H_2_O (orange) and wound + In11 (green) treatment. Independent plants were sampled at each treatment x time combination. Lines and error bars represent means ± SEM.

We also determined the expression pattern of *VuLHY* genes under constant light (LL) to confirm the presence of a free-running clock in cowpea. Both *VuLHY* genes sustained rhythmic expression for up to 48 h in LL with a peak at dusk (Fig. **4a**), although the expression level was greatly reduced after 24 h and almost abolished after 48 h. This conditional arrhythmic phenotype was also supported by the expression pattern of cowpea *GIGANTEA* (*VuGI*) homolog (Fig. **4b**) whose expression is directly regulated by CCA1/LHY1 (Lu *et al*., 2012), because it was continuously expressed at high levels throughout the day after 24 h in LL, which is consistent with lack of repression by LHY in free-running conditions. We conclude that a free-running circadian clock is dampened after 24 h in LL conditions in cowpea.

**Figure 4.**
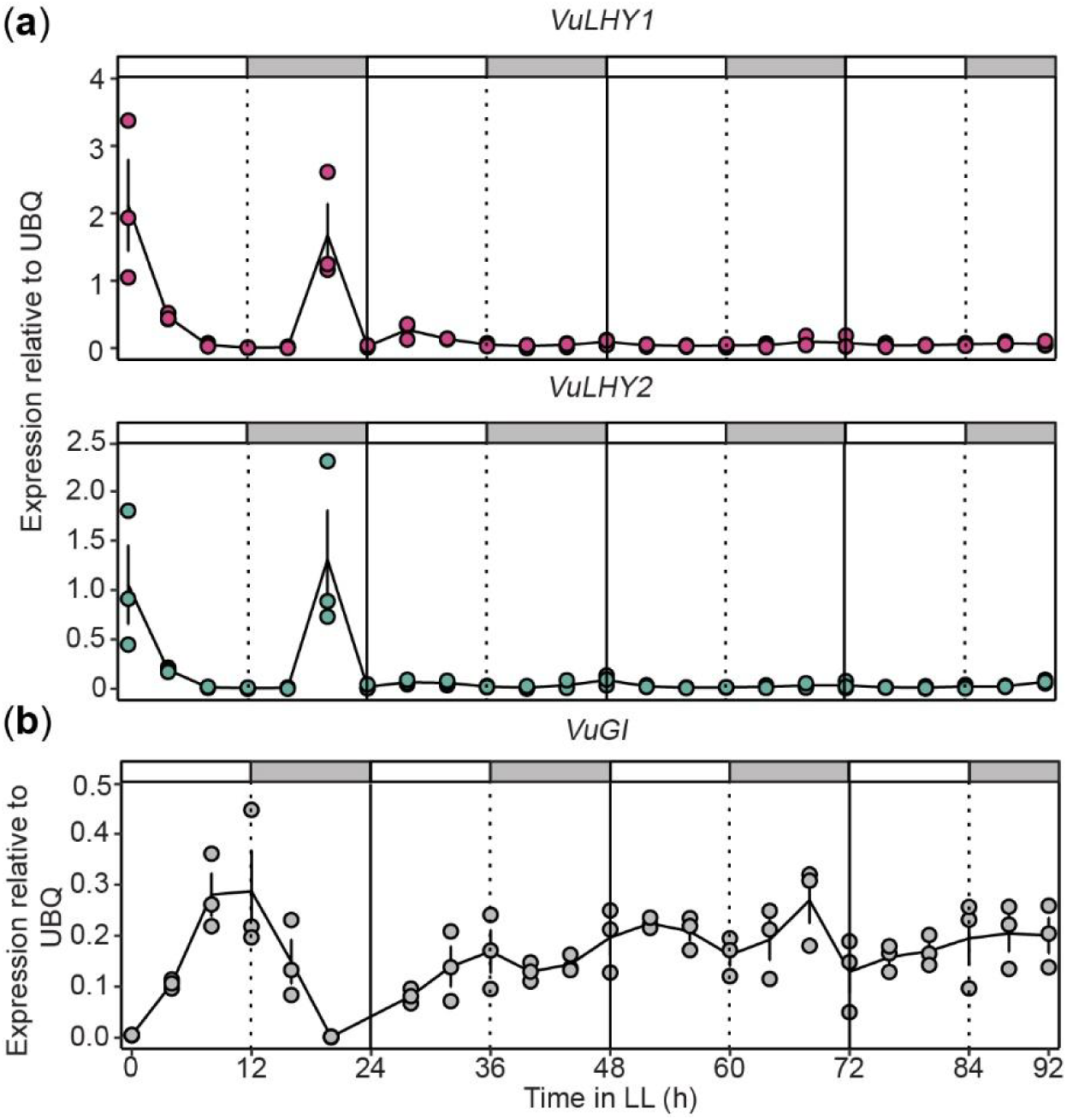
Circadian expression patterns of *VuLHY1*, *VuLHY2* and *VuGI*. Expression patterns of (a) *VuLHY1*, *VuLHY2* and (b) *VuGI* under constant light (LL) in cowpea trifoliates. Cowpea plants were grown under LD for 10 days and then transferred to LL. Leaf samples were taken every 4 hours over the course of four days for gene expression analyses. Lines and error bars represent means ± SEM (n = 3 biological replicates). Independent plants were sampled at each time point.

### The cowpea circadian clock restricts nighttime expression of In11-induced direct defenses

We used the conditional arrhythmic phenotype of cowpea plants under free-running conditions to test if nighttime repression of an In11-induced *VuKTI* was dependent on the circadian clock. We expected the time-of-day differences in In11-induced expression to be lost after 48 h in LL conditions due to reduced expression of the *VuLHY* homologs, and therefore lack of repression at nighttime. Briefly, we measured In11-induced *VuKTI* expression 6 h after treatment in plants with 4 to 88 h of LL exposure (Fig. **5a**). Consistent with our previous observations, *VuKTI* was significantly induced by In11 in subjective daytime, but not in subjective nighttime after 24 h day in LL conditions. However after 48 h in LL conditions, *VuKTI* induction by In11 was not significantly different in subjective daytime or nighttime conditions (Fig. **5b**). These results suggest that circadian oscillation of *VuLHY* is required for gated expression of In11-induced defenses to suppress *VuKTI* expression at relative nighttime when herbivore attack is less likely to happen.

**Figure 5.**
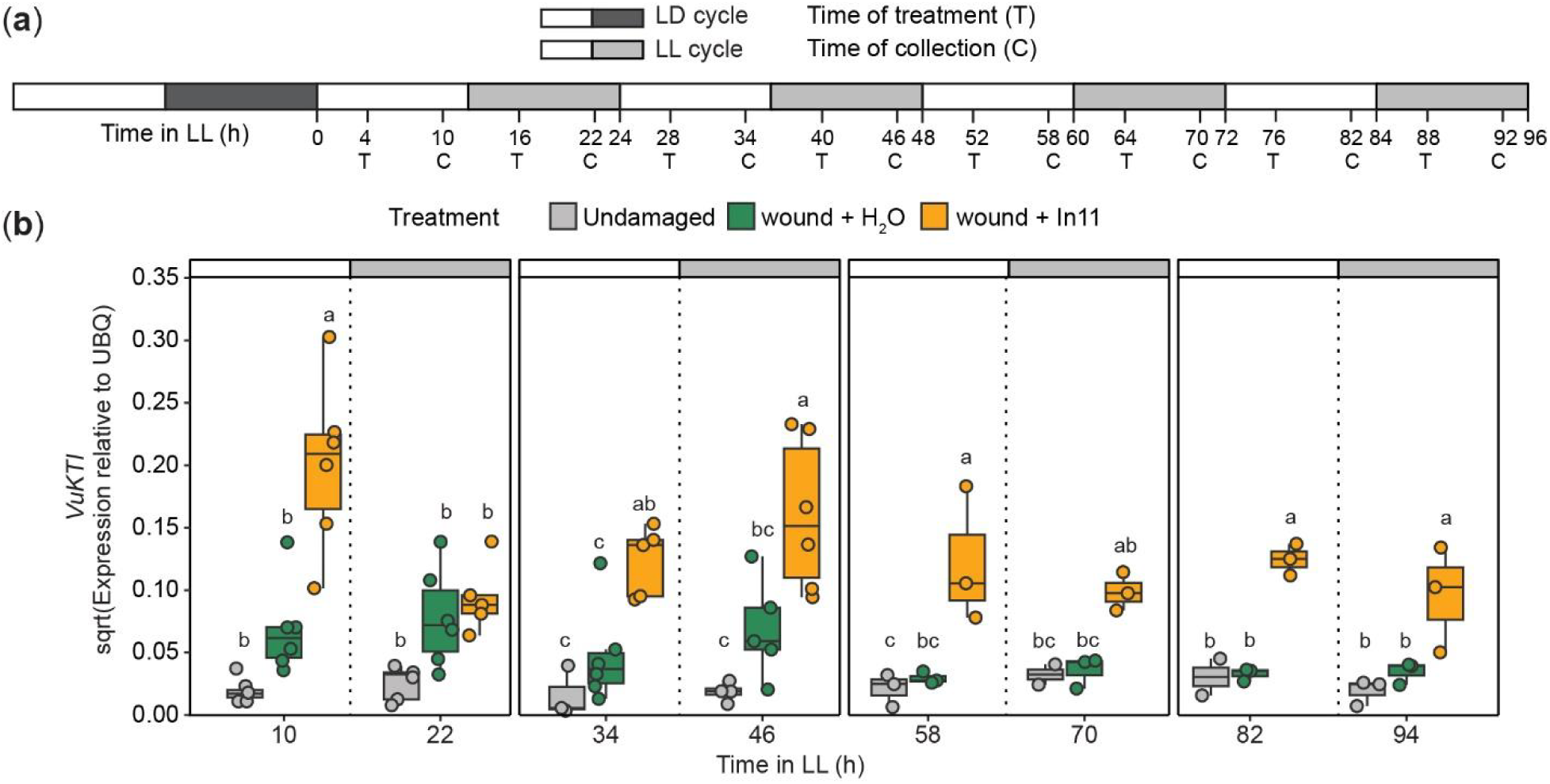
Nighttime repression of In11-induced *VuKTI* is abolished in conditional arrhythmic cowpea plants. (a) Experimental design. Cowpea plants were grown under light/dark (LD) for 10-13 days, and then transferred to constant light (LL) for one to four days. Plants were treated (T) by wound + H_2_O (orange) or wound + In11 (green) 4 h after subjective dawn or subjective dusk, and samples were collected (C) 6 h later along with undamaged (gray) controls. (b) Expression pattern of *VuKTI* according to qPCR data. Different letters indicate significant differences determined by two-way ANOVA followed by Tukey’s Honest Significant Difference test (HSD) (n= 3-6 biological replicates, p-value < 0.05) each day. Independent plants were sampled at each treatment - time combination.

### VuLHY homologs regulate *pKTI* promoter activity in a CBS-dependent manner

To test if regulation of VuKTI depends on canonical LHY-bound cis elements, we performed a transient luciferase reporter assay in *N. benthamiana* leaves. We tested both WT promoter and promoters with mutant sequences of a CBS located at position - 334 (mutant sequence M1), as well as a CBS-like (CBS-L, CAAAATCT) sequence identified at position -80 (mutant sequence M2), upstream of the TATA box (Fig. **6a,b**). Compared to the empty vector (EV) control, over-expression of VuLHY1 and VuLHY2 proteins significantly increased the activity of the *pKTI* reporter, but not when M2 was mutated. Mutations to the ATCT sequence in CBS (M1) or CBS-L (M2) resulted in less activation of the reporter by either transcription factor (Fig. **6c**). A similar transcriptional activation was also observed when *AtLHY* was overexpressed (Fig. **S4**). This data indicates that VuLHY1 and VuLHY2 interact with the In11-induced *KTI* promoter in planta via canonical and polymorphic CBSs, and that cowpea and Arabidopsis LHY homologs behave as activators when transiently overexpressed in tobacco in this context.

**Figure 6.**
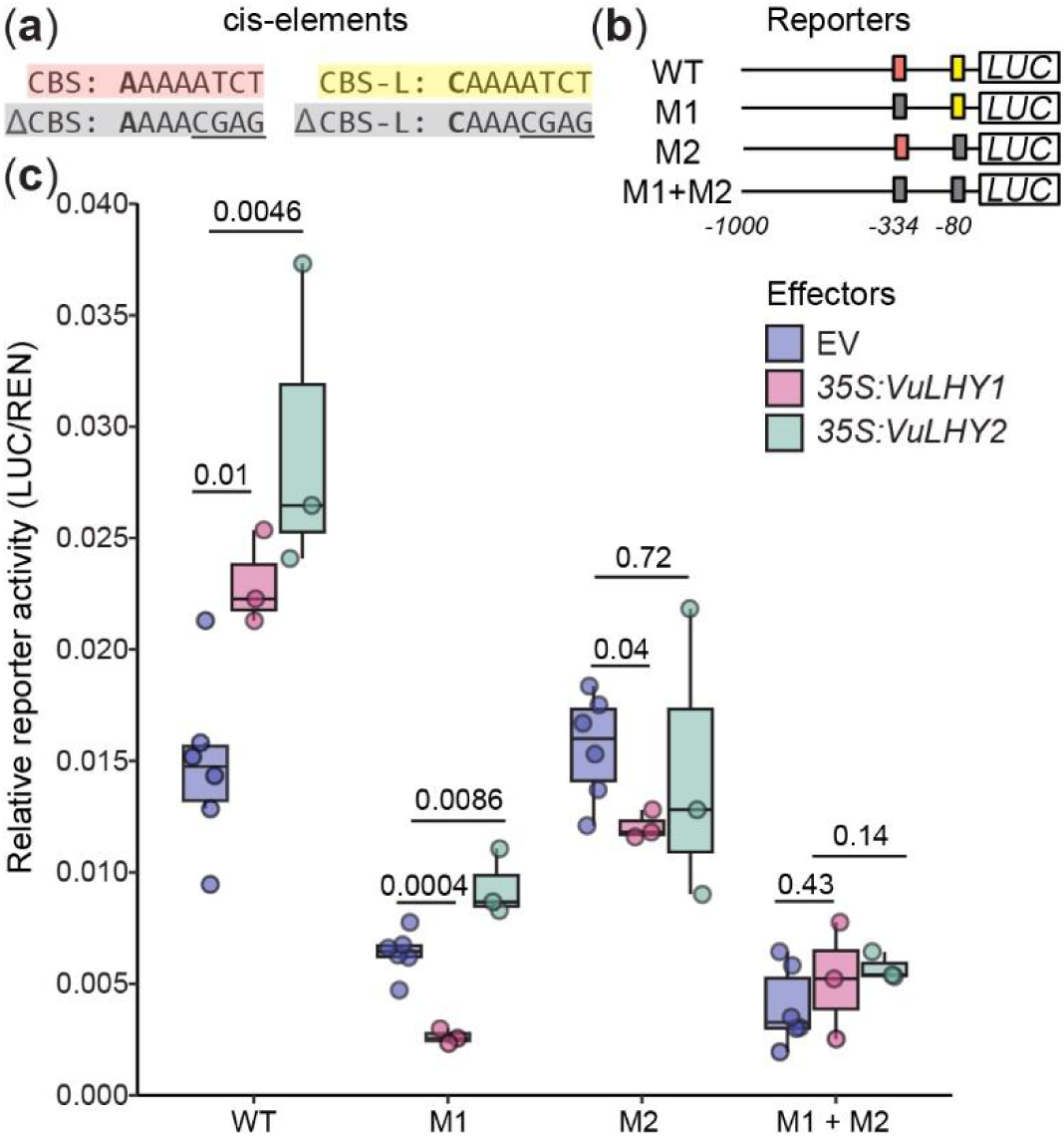
Cowpea LHY homologs modulate the activity of the *VuKTI* promoter in a CBS-dependent manner in tobacco. Schematic representation of the firefly luciferase (*LUC*) reporters used in the *N. benthamiana* transient assay (a) Sequence of the CBS and CBS-like (CBS-L) cis-elements found in the KTI promoter. The CCA1/LHY binding site was mutated on CBS and CBS-L via site directed mutagenesis (underlined) (b) LUC reporters used in the assay. WT=CBS, CBS-L, M1 = ΔCBS, CBS-L, M2 = CBS, ΔCBS-L (c) The effect of the VuLHY1 and VuLHY2 proteins on the activity of the LUC reporters. At 72 h LUC activity was measured with 35S:LHY proteins co-expressed in a separate agrobacterium strain. Relative reporter activity was calculated by normalization against *35S:Renilla*. Reporters final OD_600_=0.3 and effectors final OD_600_=0.4 Significant differences in the mean (*) were determined by a two-sided t-test of each effector vs. EV (n = 3-6 biological replicates, p-value < 0.05).

## Discussion

The time-of-day dependent modulation of transcriptional responses to specific elicitors such as HAMPs underscores the relevance of temporal cues to optimize defensive responses against herbivores. Our study of the transcriptional response of cowpea plants to HAMP In11 revealed a clear time-of-day dependence of induced gene expression associated with direct and indirect herbivore defenses and provided a mechanistic role for the plant circadian clock in directly modulating such dependence.

We found that 6 h induced gene expression was specific and significantly stronger after daytime than after nighttime elicitation with HAMP In11. The number of DEGs at ZT10 was ∼40 times larger than at night, and 50% of the GO term categories were uniquely enriched at ZT10, most of which included genes involved in antiherbivore defense like enzymes involved in volatile biosynthesis and protease inhibitors. This pattern of gene expression was not observed by wounding alone, and thus we propose that timely HAMP-specific expression of antiherbivore-related genes is tightly controlled. These findings expand our knowledge of plant defense against herbivores beyond the anticipation of attack via rhythmic accumulation of defensive hormones (Goodspeed *et al*., 2012), rhythmic accumulation of green leaf volatile (GLV) biosynthetic enzymes transcripts (Joo *et al*., 2019b), the time-of-day dependent accumulation of GLVs in response to the mix of HAMPs and effectors in caterpillar regurgitant (Joo *et al*., 2019a), and accumulation of plant volatiles in response to nocturnal and diurnal continuous mechanical damage (Arimura *et al*., 2008).

Owing to the central role of the circadian clock in regulating plant metabolism, we hypothesized that the time-of-day differences in response to In11 were in part due to direct regulation of gene expression by LHY. In support of this hypothesis, our genome-wide promoter analysis in cowpea found canonical CBS and EE located between -250 and -1000 bp upstream the coding region of any given gene (Fig. **S3b**), among which was a previously characterized VuGI homolog (Weiss *et al*., 2018), as well as a subset of In11-responsive genes with strong daytime vs nighttime differences (Fig. **2a**, Table S5). By leveraging the presence of the cis elements and a strong daytime vs nighttime difference in expression we identified multiple *Terpene synthases* (*TPS*), *chalcone synthases* (*CHS*), *chitinases*, *-glucosidases* and *Kunitz Trypsin Inhibitors* (*KTIs)* as strong candidate targets of direct regulation by VuLHY. Furthermore, we identified VuLHY1 and VuLHY2, two homologs with a conserved Myb-like DNA binding domain (Fig. **S4**) and a diurnal expression pattern that peaked at dawn under LD and LL consistent with other homologs, and thus we propose that this regulatory module is conserved in cowpea. Our RNAseq and independent qPCR data also demonstrated that wounding alone was sufficient to cause misexpression of *VuLHY1* and *VuLHY2*, and that the HAMP In11 had no further effect (Fig. 3). This indicated that abiotic stress rather than herbivory might attenuate circadian clock function, similar to damping of the circadian oscillation induced by the feedback regulation by hormones, bacterial infection, bacterial PAMPs and toxins, and unidentified molecules found in herbivore regurgitant (Zhang *et al*., 2013; Li *et al*., 2018; Joo *et al*., 2019b; de Leone *et al*., 2020; Gao *et al*., 2020; Liang *et al*., 2024b; Fraser *et al*., 2024).

Upstream factors such as the inducibility of defense hormones could explain time-of-day dependent responses, which should be apparent from transcriptional signatures of hormone biosynthesis. If hormones control strong daytime responses compared to nighttime we would expect In11-induced (Steinbrenner *et al*., 2022) biosynthetic genes such as *Allene oxide synthase* (AOS), *Allene oxide cyclase* (AOC), *Lipoxygenase* (*LOX*) for JA, and *1-aminocyclopropane-1-carboxylic* (ACC) *synthase* (*ACS*) and *ACC oxidase* (*ACO*) for ethylene, to show strong time of day dependent expression. Surprisingly, although one ACS and three LOXs are induced by In11, only VuLOX2 (Vigun11g163500) had at least one CBS element and weak daytime vs nighttime differences (ZT10-ZT22 LFC_diff_ = 0.74), indicating that In11-induced accumulation of JA in the morning might only be a small factor contributing to the enhanced daytime response to In11. Further studies of JA dynamics in time-of-day dependent In11 responses will clarify this pattern.

Using a classical free-running conditions experiment under constant light; we demonstrated that the time-of-day dependent In11 induced expression of an anti-herbivore *VuKTI* is dependent on the circadian clock. We characterized the expression of *VuLHY1*, *VuLHY2* and *VuGI* under LL and demonstrated that they oscillated only for 24 h, and became arrhythmic after; this timing pattern was similar to that of the clock of petunia leaves under DD (Fenske *et al*., 2015).

Since genetic resources and transformation methods in cowpea are lacking, the characterization of free-running conditions provides a method to study circadian regulation in emerging model systems, including other legumes and crop species

This unique circadian characteristic provided the conditional arrhythmic conditions in cowpea that later served our experiments in two ways: 1) the first 24 h after transfer to LL allowed us to address the role of light in the time-of day dependent response to In11 and 2) the following 24 to 96 h served as the conditional *VuLHY* knockdown (or arrhythmic clock) mutant. We leveraged the conditional arrhythmic plants and demonstrated that In11-induced *VuKTI* expression was higher after daytime treatment than nighttime under constant light conditions in the first 24 h of LL conditions, but that *VuKTI* was equally induced at subjective daytime and nighttime conditions once *VuLHY1* and *VuLHY2* became mis expressed (Fig. 5). We conclude that light is not a mechanism regulating morning In11-induced *VuKTI* expression, although light does partially regulate herbivore-induced terpene synthesis and emission (Arimura *et al*., 2008; Joo *et al*., 2019a), and many DEGs with strong daytime and nighttime differences did not have a canonical CBS or EE site in their promoter, indicating that light and indirect regulation by the circadian clock contribute to the overall time-of-day dependent response to In11. While direct mechanisms of regulation are difficult to study in cowpea due to lack of genetic tools, these patterns are consistent with a model where Lack of In11-induced expression of *VuKTI* during the night is likely due to transcriptional repression by the VuLHY homologs.

Our transient luciferase reporter assay demonstrated that overexpression of VuLHY1 and VuLHY2 modulated the activity of the *VuKTI* promoter in a CBS dependent manner. By comparing the activity of reporters bearing wild type and mutated variants of the CBS sites, we determined that changes to the ATCT sequence in the 5’ end of the element are sufficient to alter the interaction between the promoter and the transcription factor. This is similar to the interaction of AtLHY with CBS and EE (Harmer *et al*., 2000; Nagel *et al*., 2015; Kamioka *et al*., 2016; Adams *et al*., 2018; Kim *et al*., 2023) via this sequence (Wang *et al*., 1997), further supporting that VuLHY targets genes via CBS. We also found a polymorphic variant that we have named CBS-like (CBS-L: CAAAATCT) that also requires a conserved 5’ end to interact with both VuLHY homologs. Based on the distribution and abundance of CBS-L (Table S6), we propose it is likely a novel VuLHY binding site in cowpea. Consistent with our results, Arabidopsis LHY binds other sequences in genome-wide analyses (Adams *et al*., 2018). The higher background level activity of CBS-L co-expressed with the EV, and the differential effect on interaction with VuLHY1 and VuLHY2 suggests that this site might provide some specificity of binding and an added layer of regulation under certain conditions, although this remains to be explored in detail. In our transient system both VuLHY1 and VuLHY2 functioned as activators, likely due to the regulatory environment in the *N. benthamiana* transient expression system since AtLHY, typically a repressor, also functioned as a weak activator under our experimental conditions (Fig. **S4a**), although a unique activation function has been described for AtLHY in the fatty acid synthesis pathway (Kim *et al*., 2023). Nevertheless, our free running experiment using the conditional arrhythmic plants clearly demonstrated that *VuGI,* a possible direct target, became arrhythmic and highly expressed when *VuLHY* expression was low, thus supporting a repressive function for *VuLHY* against its regulated target genes.

In summary, we describe a molecular link between the plant circadian clock and HAMP-induced gene expression in cowpea. VuLHY gates the expression of In11-induced genes likely fine tuning the herbivore-specific response. At night when *VuLHY* is highly expressed, VuLHY interacts with the promoter of In11-responsive genes involved in antiherbivore defense such as *VuKTI* to repress their expression. When *VuLHY* expression decreases during daytime, the CBS-bearing In11-induced promoters are available for recruitment of the transcriptional machinery required for antiherbivore response. The relevance of this regulation to physiology and metabolism of anti-herbivore defenses is a topic that should be further explored. We expect that gating of HAMP-induced responses by the circadian clock is a mechanism to minimize the effect of the growth-immunity trade-off by allowing robust and specific response during the day without interfering with nighttime growth.

## Acknowledgments

This research is supported by NIH 5R35GM151272 and NSF 2139986 to ADS and NIH R01GM079712 to TI. We thank members of the Nemhauser and Di Stilio labs for conversation and feedback. NGP and ADS were supported by start-up funding from the University of Washington. NGP is supported by the USDA-AFRI predoctoral fellowship Grant #2023-67011-40362 and was partially supported by the UW Royalty Research Fund grant #A161929 and the Hereensperger and Walter and Margaret Sargent Awards.

## Author Contributions

N.G.P., T.I., A.D.S. conceived and designed the study. N.G.P. performed the experiments, analyzed the data, and wrote the article. All the authors critically read and commented on the article and approved its final version for submission.

## Data availability

All transcriptomics data is available at the National Center for Biotechnology Information (NCBI) under Bioproject PRJNA1168576.

## Supporting information

**Table S1.** List of primers used in this study.

**Table S2.** FIMO summary of the AAMWATCT motif for all promoters in the cowpea genome.

**Table S3.** List of early (1 h) and late (6 h) DEGs after daytime and nighttime w + H_2_O or w + In11 treatment.

**Table S4.** Significantly enriched (padj < 0.05) Gene Ontology (GO) categories among early (1 h) and late (6 h) w + In11 DEGs.

**Table S5.** Early (1 h) and late (6 h) w +In11 DEGs and their LFCdiff daytime-nighttime and CBS/EE counts.

**Table S6.** FIMO summary of the CBS-L (CAAAATCT) site for all promoters in the cowpea genome.

**Figure S1.**
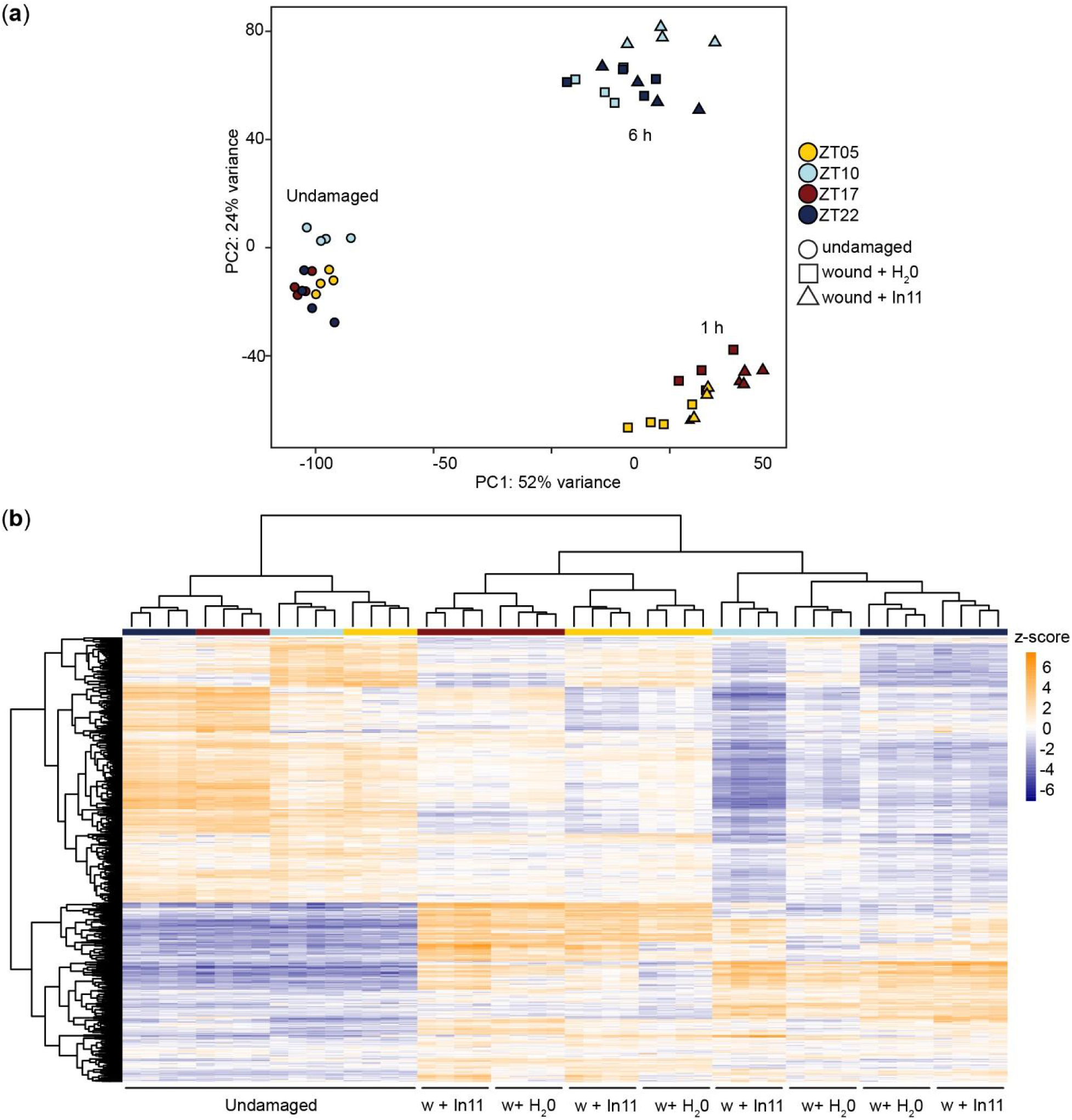
Clustering analysis of DEGs in response to w + H_2_O and w + In11. (a) Principal component (PC) analysis of differentially expressed genes (DEGs) across all samples. (b) Hierarchical clustering of samples according to the expression pattern of 847 In11-responsive DEGs across all samples.

**Figure S2.**
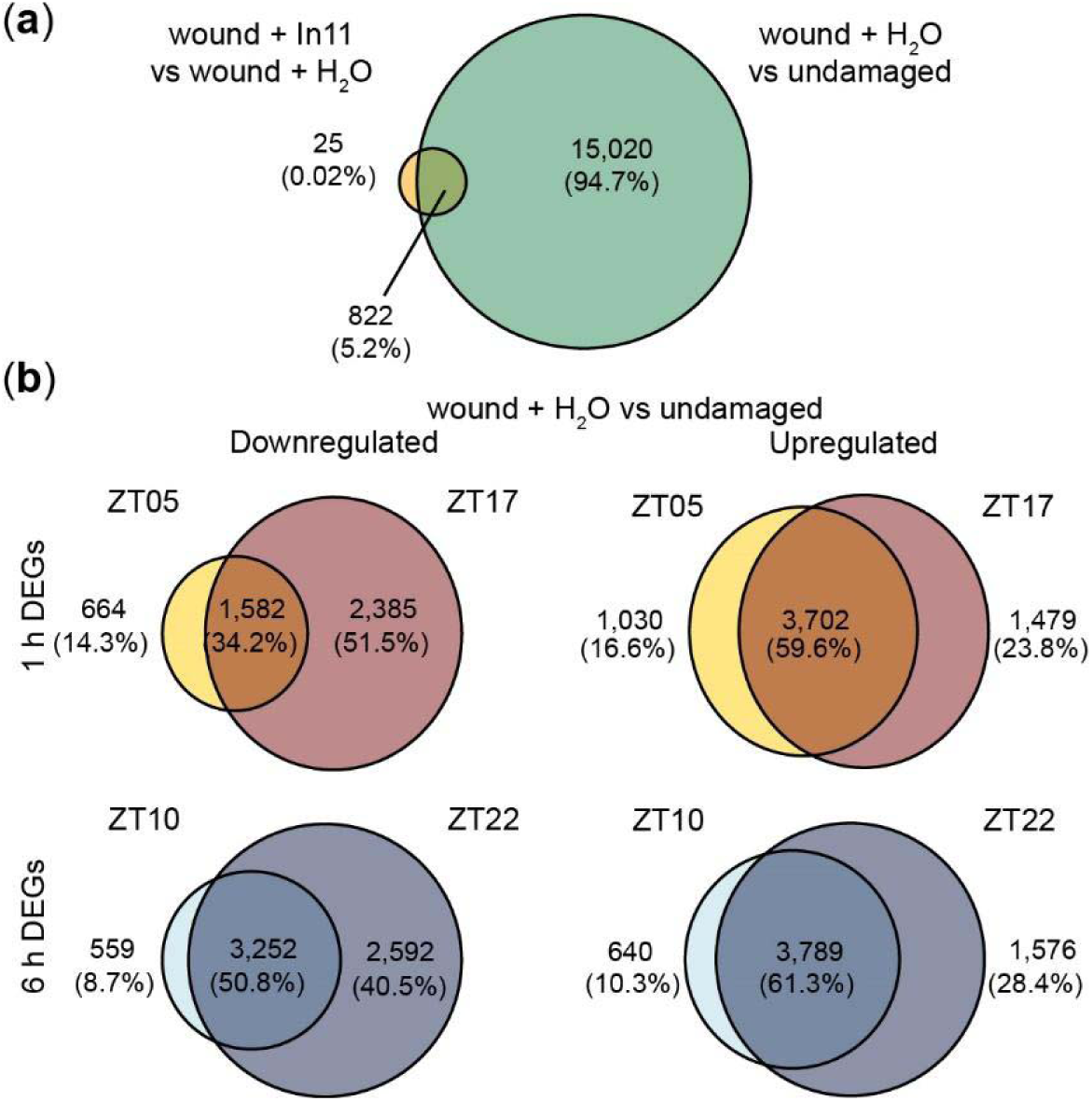
Time -of-day response to wound + H_2_O vs undamaged. Venn diagrams indicating the number of shared and unique up and down-regulated genes (a) In11 vs wound across all time points, and (b) 1h (ZT5 vs ZT17) and 6h (ZT10 vs ZT22) after daytime or nighttime wounding (wound + H_2_O vs undamaged).

**Figure S3.**
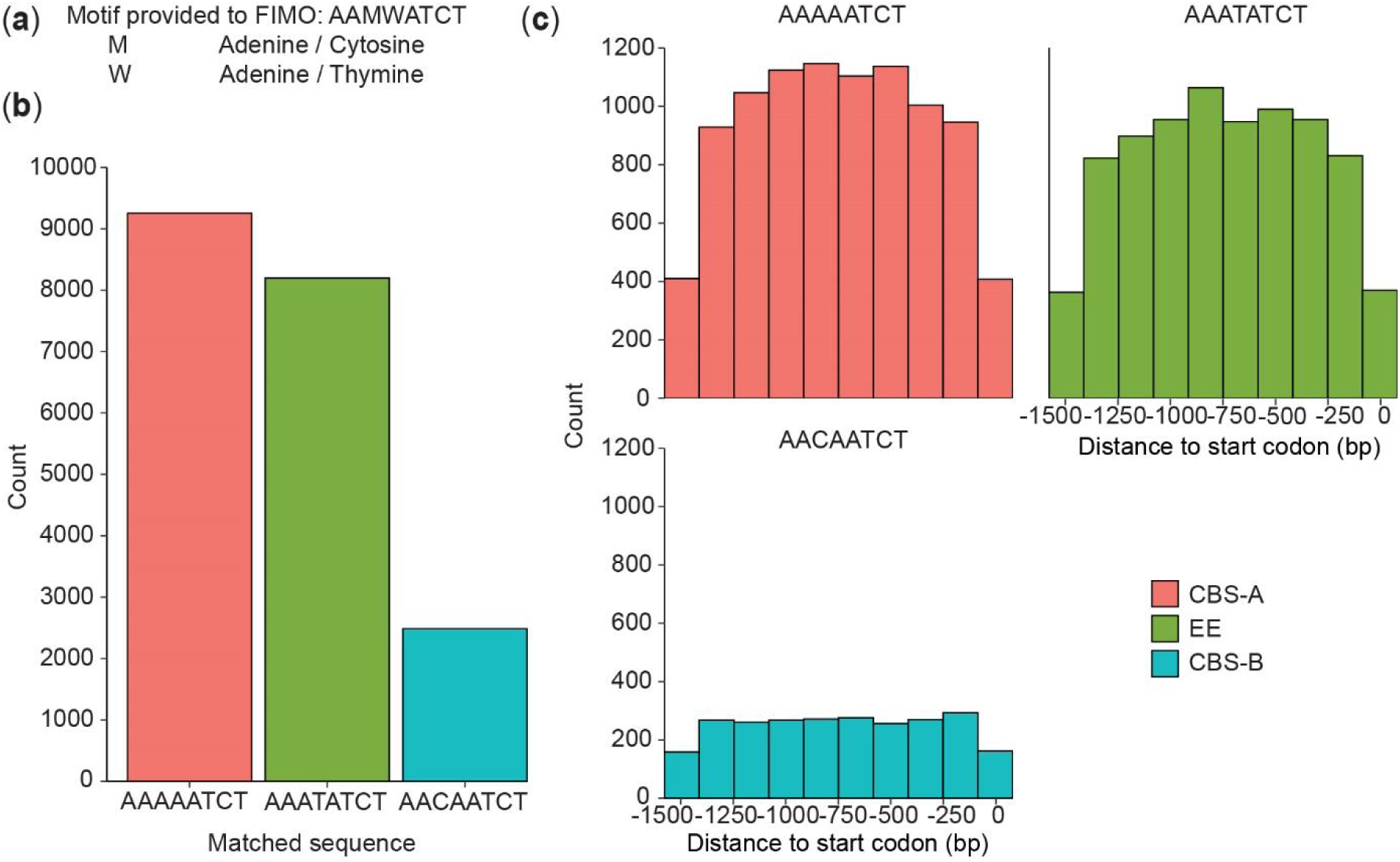
Genome-wide distribution and abundance of CBS and EE motifs in cowpea promoters. Predicted promoter sequences (1.5 kb upstream start codon) were retrieved from the cowpea genome for a circadian clock cis element analysis. (a) Motif provided to Find Individual Motif Occurrences (FIMO) software. (b) Sequence and count for known and novel motifs found in the promoters. CCA1 Binding site A (CBS-A), Evening element (EE), CBS-B, CBS-like. (c) Motif location distribution in the promoters in 250 base pair (bp) bins.

**Figure S4.**
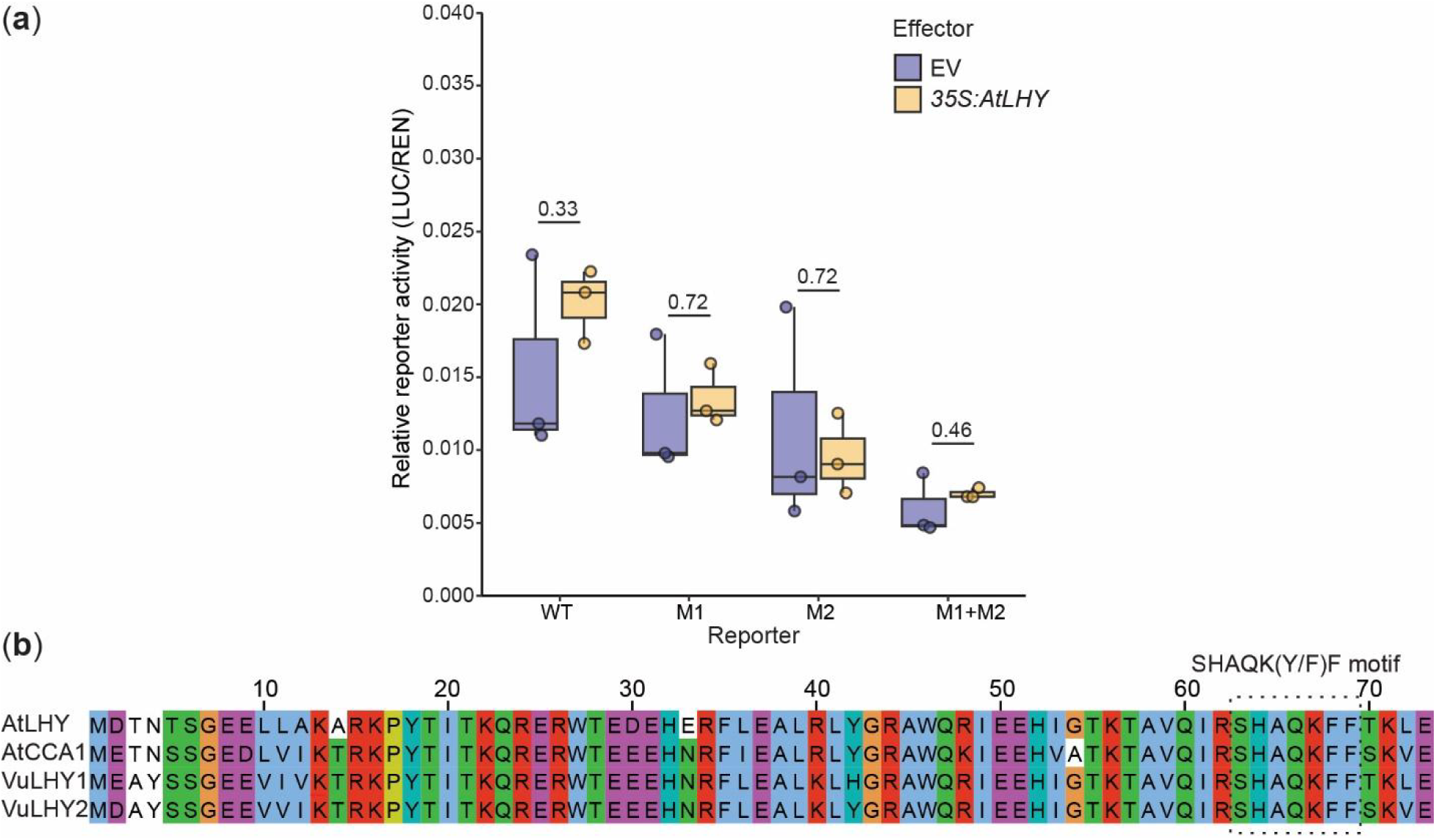
AtLHY weakly activates the *VuKTI* promoter in a CBS-dependent manner. The effect of the AtLHY protein on the activity of the LUC reporters. (a) At 72 h LUC activity was measured with 35S:AtLHY protein co-expressed in a separate agrobacterium strain. WT=CBS, CBS-L, M1 = ΔCBS, CBS-L, M2 = CBS, ΔCBS-L. Relative reporter activity was calculated by normalization against *35S:Renilla*. Reporters final OD_600_=0.3 and effectors final OD_600_=0.4 Significant differences in the mean (*) were determined by a two-sided t-test of each effector vs. EV (n = 3, α= 0.05). (b) Alignment of the amino acid sequence of the Myb-like DNA binding domain for AtLHY, AtCCA1, VuLHY1 and VuLHY2. Color scale according to ClustalW.

